# Experience in developing the human genome standard E701

**DOI:** 10.1101/2024.09.30.615836

**Authors:** Iuliia Vasiliadis, Vera Belova, Anna Shmitko, Anna Kuznetsova, Alina Samitova, Oleg Suchalko, Andrey Goltsov, Peter Shatalov, Peter Shegai, Olga Melkova, Tatiana Kulyabina, Elena Kulyabina, Denis Rebrikov, Dmitriy Korostin

## Abstract

The first Russian human genome standard E701 was developed through a collaborative research involving four laboratories: Pirogov Russian National Research Medical University, National Medical Research Center for Obstetrics, Gynecology and Perinatology named after Academician V.I.Kulakov, National Research Center Kurchatov Institute, and National Medical Research Radiological Centre. Whole-genome sequencing of short reads on various platforms (MGI Tech, Illumina) and alignment to the reference human genome GRCh38.p14 were performed for the E701 sample. Subsequently, 3842877 high confidence genomic variants were identified, which can be used as a standard for calculating statistical quality metrics while analyzing sequencing data. Furthermore, 9096 biallelic variants were identified on the autosomes and the X chromosome, with a minor allele frequency exceeding 0.4. Additionally, mitochondrial DNA sequencing was performed with the breadth of coverage over 99.9% at 1000X.

## Introduction

The rapid development of next-generation sequencing (NGS) technologies imposes new demands on the reliability of identified genetic variants and the potential for errors in bioinformatic data analysis. Research shows that results obtained from different sequencing platforms and bioinformatics algorithms can vary significantly [1,2,3,4,5]. A highly accurate reference is essential for evaluating the accuracy and reproducibility of various sequencing methods. The most well-known human reference genomes were developed by the Genome in a Bottle (GIAB) Consortium and are widely used for sequencing validation [6,7].

Using high-throughput sequencing from multiple platforms and bioinformatic analysis of the obtained data, we produced the first Russian human genome standard E701.

## Materials and methods

### Ethics Statement

This study conformed to the principles of the Declaration of Helsinki. The appropriate institutional review board approval for this study was obtained from the Ethics Committee at the Pirogov Russian National Research Medical University. Patient provided written informed consent for sample collection, subsequent analysis, and publication thereof.

### DNA source

The genomic DNA derived from peripheral blood mononuclear cells came from a white Caucasian male with karyotype 46XY with no history of hereditary pathologies. Whole blood was collected in EDTA-Vacutainer tubes and the DNA extraction was performed using the QIAamp DNA Blood Kit (Qiagen) according to the manufacturer’s protocol.

### Sequencing

Whole-genome sequencing (WGS) data in paired-end (PE) mode with 30X coverage were obtained using various high-throughput sequencing platforms in four laboratories: Pirogov Russian National Research Medical University (RSMU), National Medical Research Center for Obstetrics, Gynecology and Perinatology named after Academician V.I.Kulakov (NCAGP), National Research Center Kurchatov Institute (NRCKI), and National Medical Research Radiological Centre (NMICR) (Table 1). Library preparation followed the manufacturer’s instructions (MGI and Illumina).

**Table 1.**
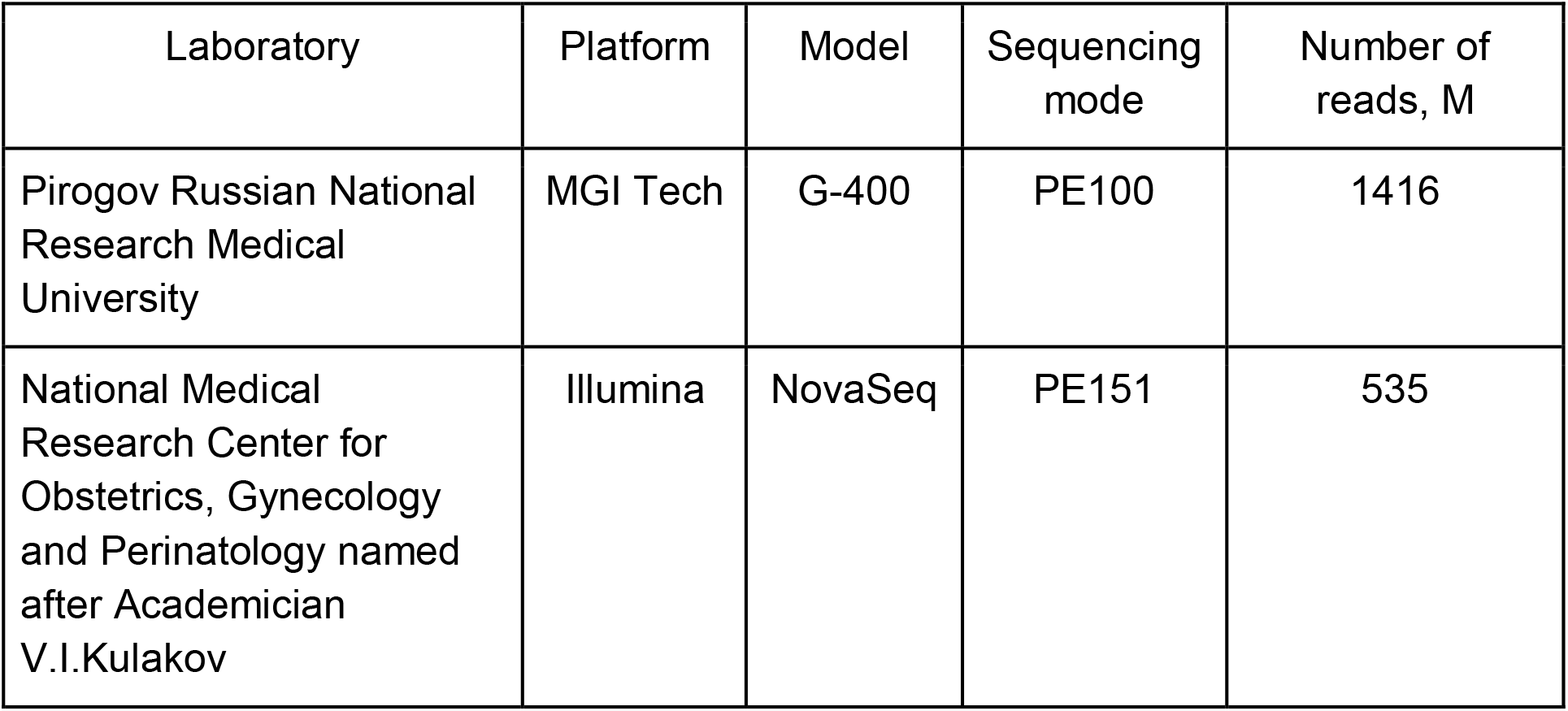
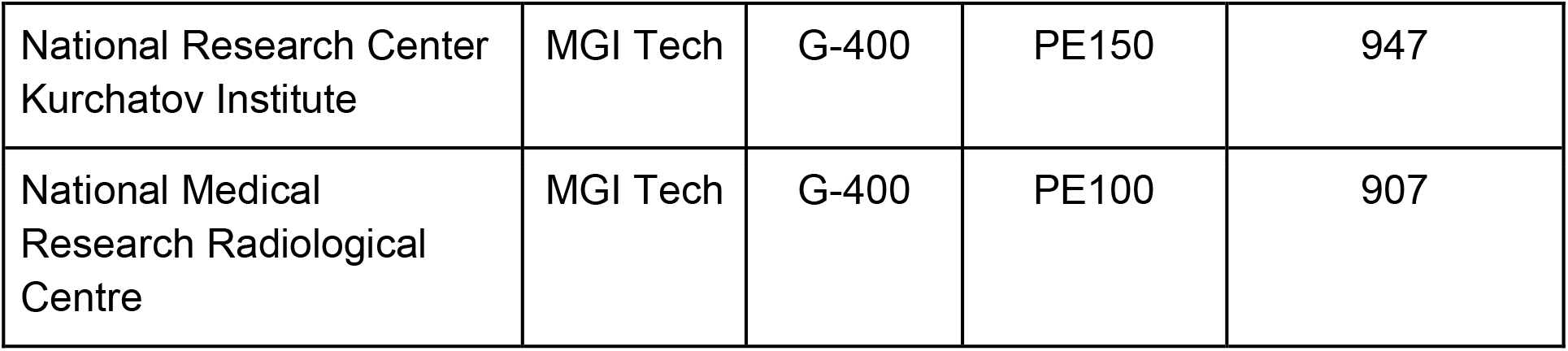
Description of sequencing platforms and raw data quantity.

### Bioinformatics processing

The quality of the obtained paired FastQ files was analyzed using FastQC v0.11.9 [8]. Based on the quality metrics, the FastQ files were trimmed using BBDuk by BBMap v38.96 [9]. Reads were aligned to the indexed reference genome GRCh38.p14 (https://www.ncbi.nlm.nih.gov/datasets/genome/GCF_000001405.40/) using bwa-mem2 v2.2.1 [10]. SAM files were converted into BAM files and sorted using SAMtools v1.9 [11]. The number of duplicates was calculated using Picard MarkDuplicates v2.22.4 [12].

## Results

### Efficiency of whole-genome sequencing

To assess the alignment quality and uniformity of genome coverage, Picard CollectWgsMetrics and SAMtools flagstat were used. The results showed that over 99% of reads were aligned to the reference genome, indicating the high quality of the raw data and the efficiency of the alignment algorithm used. Median coverage of the sample obtained from each laboratory was at least 20 reads. Mean coverage for WGS E701 was 39X in the RSMU sample, 32X in the NRCKI sample, 24X in the NMICR sample, and 19X in the NCAGP sample. For all four samples, the breadth of coverage at 1X was approximately 91% and 5X about 90%. The breadth of coverage at 10X varied slightly across samples with a maximum value of 89.96% observed in the RSMU sample and a minimum value of 86.94% observed in the NCAGP sample. The values for the other metrics are presented in Supplementary table 1.

### Variant calling

Variant calling was performed using two programs: bcftools mpileup v1.9 [13] and DeepVariant v1.5.0 [14]. Subsequently, multiallelic variants in VCF files were decomposed into biallelic variants using vt decompose v0.5772 [15] and then normalized using vt normalize v0.5772. A depth coverage filter (DP>=3) was applied to variants called by both programs. An additional filter (FILTER=PASS) was applied to variants called by DeepVariant, after which VCF files were merged using bcftools-1.9 merge.

### Combined VCF

To identify reliable variants in the E701 sample, we filtered single nucleotide variants (SNVs) in each of the four VCF files using bcftools-1.9 view. Subsequently, the filtered VCF files were intersected using the bcftools-1.9 isec and custom-written scripts. The intersection yielded a list of 3842877 variants (Fig. 1), which served as a true set of SNVs, forming a benchmark variant call set. This approach significantly reduced the probability of including erroneous variants in the research results and ensured high reliability of the data obtained.

**Figure 1.**
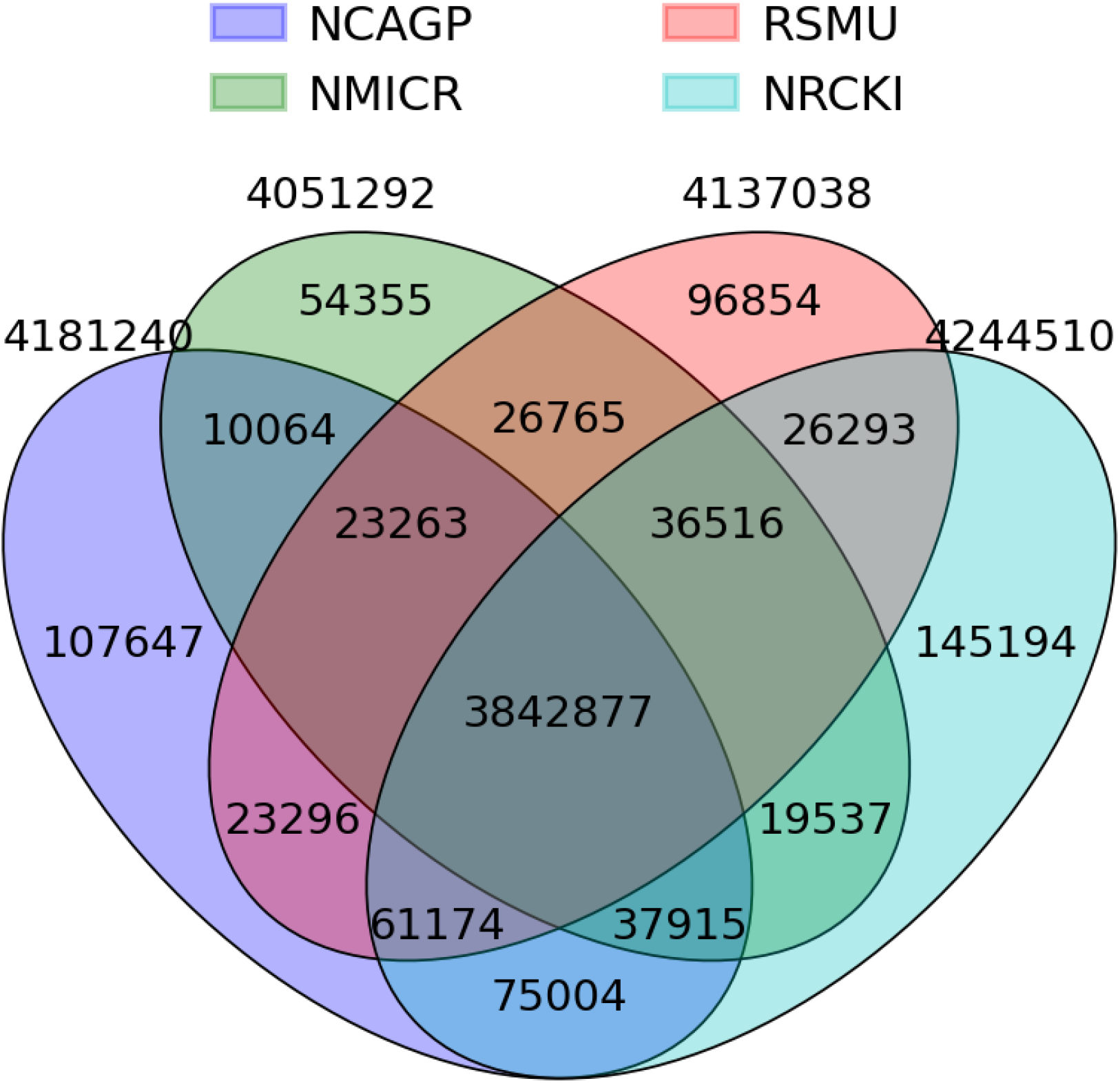
Venn diagram showing the overlap of variants in the E701 sample from four laboratories

### Assessment of variant identification accuracy

To evaluate the results of variant calling, we used the dbSNP (Database of Single Nucleotide Polymorphisms) database release 156 (https://ftp.ncbi.nlm.nih.gov/snp/archive/b156/VCF/GCF_000001405.40.gz) [16]. The multiallelic variants from the database were split into biallelic variants and then crossed with the variants called in the E701 sample. According to the results, 99.59% (3827006) of the variants identified in the E701 sample were present in dbSNP database (Fig. 2).

**Figure 2.**
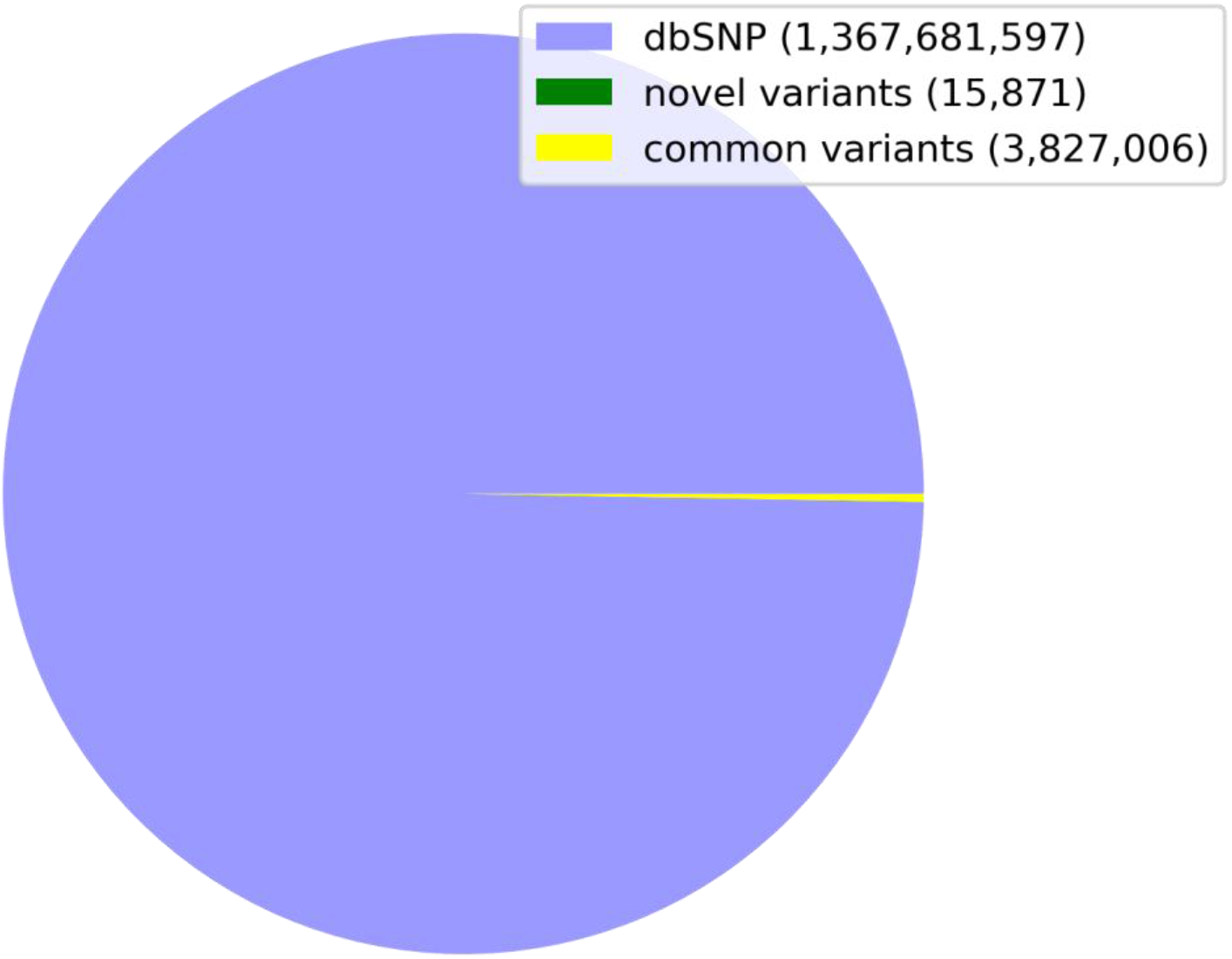
The number of detected SNPs in the E701 sample that overlap with variants in the dbSNP database

### “EXOME-GID”

Based on the data extracted from the Genome Aggregation Database (gnomAD), a list of 9096 biallelic variants was obtained. These variants are located on autosomes and the X chromosome, within coding regions of the genome, are not associated with pathogenic traits, and have a global minor allele frequency (GMAF) in the range of 0.4 to 0.5 [17].

### Ti/Tv

One of the metrics for assessing the accuracy of variant calling is the identification of point mutation types: transitions (Ti) and transversions (Tv), the ratio of which for the whole human genome is typically within the range of 2 to 2.1. The Ti/Tv ratio was calculated using the bcftools stats tool for the set of variants shared by four laboratories, yielding a value of 2.01. This result is consistent with the expected result for whole genome data.

### Mitochondrial DNA

Human mitochondrial DNA (mtDNA) is a small circular molecule of about 16.6 Kb (16,569 bp) that is maternally inherited and exhibits a higher mutation rate compared to nuclear DNA [18].

For the E701-MT sample, the breadth of coverage at 1000X (PCT_1000X) was 99.994%. The mean coverage depth was 22880 reads, with a minimum of 841 reads observed at position 3107, labeled N in the mitochondrial chromosome reference assembly. The values for other metrics are presented in Supplementary table 1.

Extraction of reads unambiguously mapped to the mitochondrial chromosome and obtaining a consensus sequence based on them was performed using SAMtools-1.9. Variant calling and annotation were conducted using mutserve [19]. The variants that differ between E701-MT sample and the mtDNA reference assembly (hg38) are listed in Supplementary table 2. Variants flagged by the “artifact_prone_site” filter were excluded from the final variant list.

## Conclusion

We present the first Russian human genome standard E701, which was developed through a collaborative effort involving four research laboratories (Pirogov Russian National Research Medical University, National Medical Research Center for Obstetrics, Gynecology and Perinatology named after Academician V.I.Kulakov, National Research Center Kurchatov Institute, and National Medical Research Radiological Centre). Whole-genome sequencing on various platforms was performed for the E701 sample, with sequencing and alignment efficiency evaluated against the human reference genome GRCh38.p14. We identified 3842877 genomic SNVs matched across the four laboratories, which can serve as a benchmark set to filter false positive and false negative variants generated during sequencing data processing. Of the variants identified, more than 99% were found in the dbSNP database. We also obtained a list of 9096 biallelic variants located in autosomes and the X chromosome, with a minor allele frequency exceeding 0.4. In addition to genomic DNA, mitochondrial DNA was sequenced with a breadth of coverage over 99.9% at 1000X and mean coverage depth of 22880X.

Future work will involve sequencing the family members of the E701 genetic material donor and integrating long-read sequencing technologies to improve the detection of indels and structural variants.

## Data availability

Genome sequences and alignments for the E701 WGS sample in fastq.gz and BAM formats, and sequences for E701-MT mitochondrial DNA in fastq.gz have been deposited into the NCBI open-access sequence read archive (SRA) under the BioProject number PRJNA1083205.

